# Sex-Specific Effects of Early Life Unpredictability on Hippocampal and Amygdala Responses to Novelty in Adolescents

**DOI:** 10.1101/2024.09.20.614130

**Authors:** Elysia Poggi Davis, Bianca T. Leonard, Robert J. Jirsaraie, David B. Keator, Steven L. Small, Curt A. Sandman, Victoria B Risbrough, Hal S. Stern, Laura M. Glynn, Michael A. Yassa, Tallie Z. Baram, Jerod M. Rasmussen

## Abstract

**Background:** Unpredictable childhood experiences are an understudied form of early life adversity that impacts neurodevelopment in a sex-specific manner. The neurobiological processes by which exposure to early-life unpredictability impacts development and vulnerability to psychopathology remain poorly understood. The present study investigates the sex-specific consequences of early-life unpredictability on the limbic network, focusing on the hippocampus and the amygdala.

**Methods:** Participants included 150 youth (54% female). Early life unpredictability was assessed using the Questionnaire of Unpredictability in Childhood (QUIC). Participants engaged in a task-fMRI scan between the ages of 8 and 17 (223 total observations) measuring BOLD responses to novel and familiar scenes.

**Results:** Exposure to early-life unpredictability associated with BOLD contrast (novel vs. familiar) in a sex-specific manner. For males, but not females, higher QUIC scores were associated with lower BOLD activation in response to novel vs. familiar stimuli in the hippocampal head and amygdala. Secondary psychophysiological interaction (PPI) analyses revealed complementary sex-specific associations between QUIC and condition-specific functional connectivity between the right and left amygdala, as well as between the right amygdala and hippocampus bilaterally.

**Conclusion:** Exposure to unpredictability in early life has persistent implications for the functional operations of limbic circuits. Importantly, consistent with emerging experimental animal and human studies, the consequences of early life unpredictability differ for males and females. Further, impacts of early-life unpredictability were independent of other risk factors including lower household income and negative life events, indicating distinct consequences of early-life unpredictability over and above more commonly studied types of early life adversity.

## Introduction

Early-life adversity (ELA) is now well established to have long-term neurodevelopmental consequences with increased risk of adverse mental health outcomes (1–3). However, an incomplete understanding of the neurobiological mechanisms linking exposure to ELA to mental health outcomes constrains the development of targeted prevention strategies. Moreover, while most ELA research has focused on the severity or frequency of adverse events, emerging evidence across species supports unpredictable patterns in the early environment as an additional, distinct, and potent form of ELA that impacts neurodevelopment in a sex-specific manner (4–11).

Unlike traditional measures of ELA that focus on the severity or frequency of adverse events, unpredictability pertains to the inconsistency and variability of environmental signals, which can uniquely disrupt developmental processes (1, 12). Given this, it is essential to consider how such variability influences children’s interactions with novel stimuli. Indeed, recent evidence suggests that the exploration of, and responses to novelty are essential for healthy neurodevelopment and are developmentally plastic in the context of unpredictability (3). Further, studies from several groups including our own, now suggests that learning, memory, and the processing of novel information are strongly susceptible to early life unpredictability (13–15). Unpredictability is also associated with increased anxiety, depression and post-traumatic stress disorder symptoms in humans independent of other forms of adversity (7, 16). Therefore, it is increasingly important to understand the underlying neural circuits and mechanisms involved.

A growing and convergent body of cross-species literature demonstrates that exposure to unpredictable parental signals in early life disrupts memory-related processes including distinguishing novel and familiar objects (5, 13, 17–19). Moreover, compelling evidence indicates that early life unpredictability alters child exploration behaviors, including exploration of novelty (15). Importantly, these behavioral phenotypes are consistent with observed associations between early life unpredictability and anxiety symptoms (20–24), thus motivating examination of responses to novelty as a construct of interest in understanding the role of early life unpredictability in the developmental origins of mental health disorders (1).

Processing of novelty engages several regions in the limbic network, most prominently the hippocampus (25) and amygdala (26). It is thus, not surprising, that these two regions are both among the most vulnerable to the effects of ELA (27–29), including unpredictability, and implicated in a host of mental health disorders including anxiety, depression, and post-traumatic stress disorder (30, 31). Notably, ELA, particularly unpredictability, exerts sex-specific consequences on the development of limbic neural circuits (11, 32, 33) and related functions (1, 9–11, 33–35) in both humans and rodents (36). For example, unpredictable patterns of maternal sensory signals associate with the integrity of the uncinate fasciculus (a white matter pathway connecting amygdala/hippocampus with the prefrontal cortex) in a sex-specific manner, affecting females but not males (19). Given these considerations, we probe here the sex-specific effects of early life unpredictability on the neural response to novel stimuli focusing on the hippocampus and amygdala.

To test the hypothesis that early-life unpredictability affects hippocampal and amygdalar responses to novelty in a sex-dependent manner, we conducted a longitudinal fMRI study based on the response to novel relative to familiar stimuli in 150 participants with a total of 223 imaging sessions across childhood and adolescence (ages 8-17 years). Early life unpredictability was characterized using the well-validated Questionnaire of Unpredictability in Childhood (QUIC). Primary analyses considered sex-specific associations between the QUIC and novel relative to familiar BOLD contrasts using mixed-effects models. This task and others like it, in the context of hippocampal/amygdala activation, has been validated in detail (37–39), and developmental changes in this cohort have been previously described (40). To test whether associations were specific to unpredictability, we conducted specificity analyses covarying for other commonly studied forms of ELA. Finally, because the amygdala and hippocampus function as part of a broader circuit, we conducted a secondary analysis to examine their context-dependent (novel vs. familiar) *functional connectivity* using a traditional psycho-physiological interaction (PPI) model.

## Methods

### Study Overview

Participants included 150 youth (54% female) participating in a longitudinal study of maternal and child health (4). Participants completed functional scanning one to three times between the ages of 8 and 17 (mean age 12.5±2.0 [S.D.]; Figure 1). The study protocol was approved by the institutional Review Boards at the University of California Irvine and Chapman University and written, and informed consent was obtained from the mothers and assent from the children. See Table 1 for demographic information.

**Figure 1.**
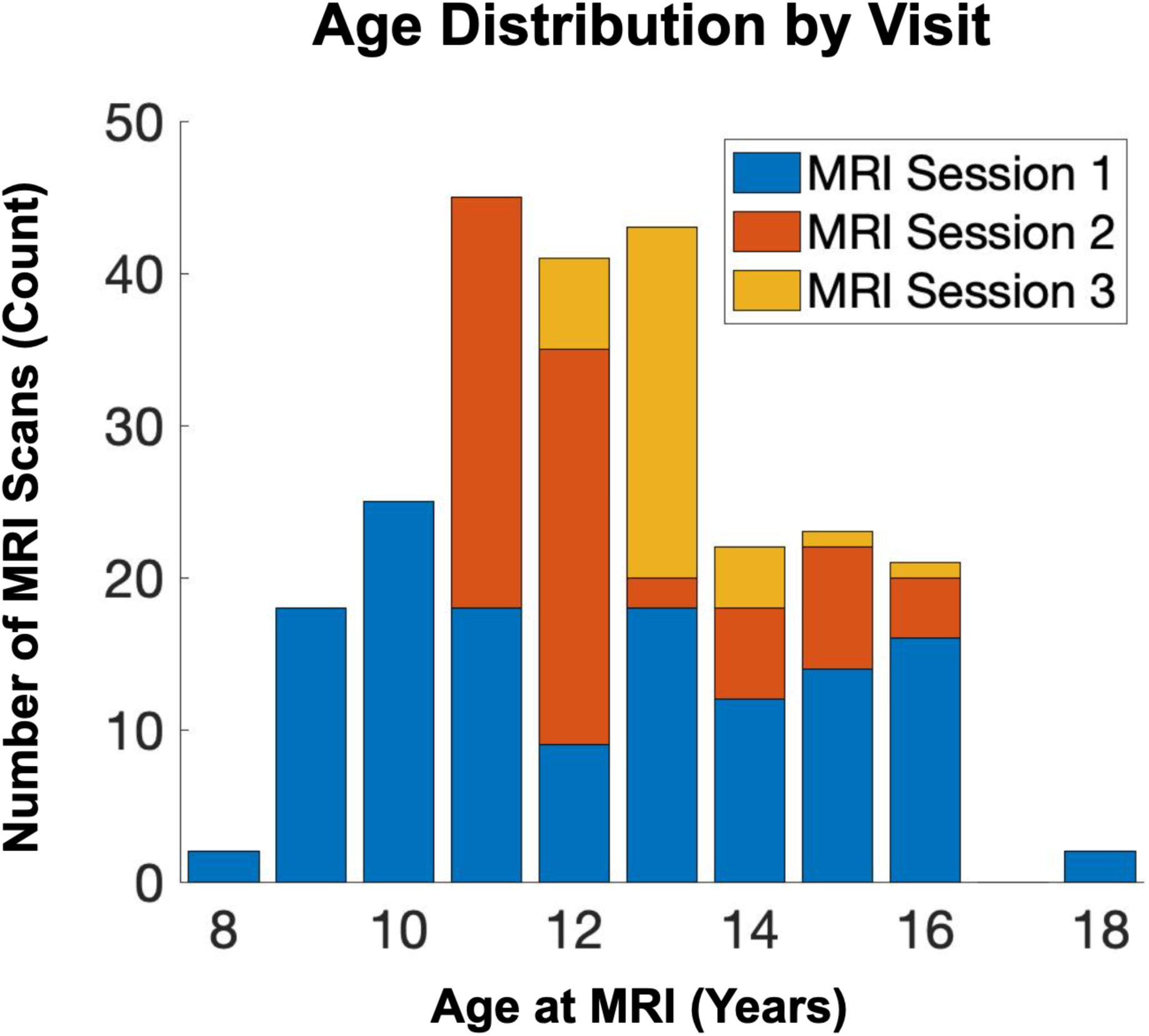
Age Distribution by Visit. Age distribution across imaging sessions/visits is shown.

**Table 1.**
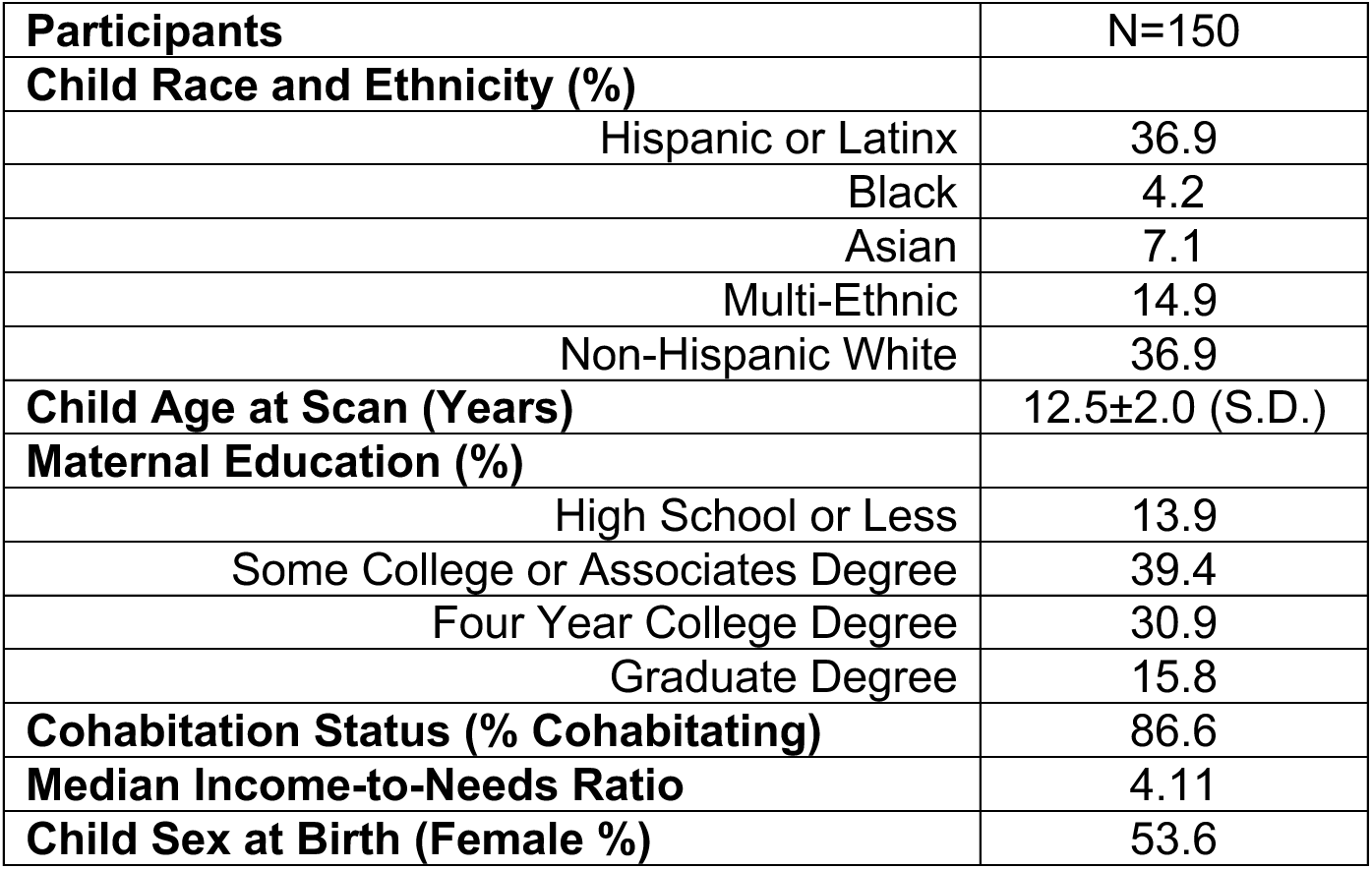
Participant Demographics Characteristics.

### Measurement of Early Life Unpredictability

Early life unpredictability was characterized using the validated 38-item version of the Questionnaire of Unpredictability in Childhood (QUIC) (22). In brief, the QUIC includes a series of items assessing unpredictability in the social, emotional and physical domains that participants endorse as applying to their life prior to the age of 18, or their current age if less than 18. For example, “I experienced changes in my custody arrangement”, and “At least one of my parents regularly checked that I did my homework”. QUIC scores range from 0 to 38, with a higher score indicating greater exposure to unpredictability. QUIC scores used here were based on self-report and acquired at the first visit (mean age 12.2+/-1.3 years old). Notably, the QUIC shows strong internal consistency (α = .89) and test-retest reliability (*r* = .92) (22). The QUIC also demonstrates convergent validity with prospective, observational measures of unpredictability in infancy, such as unpredictable maternal sensory signals (*r* = .23) and unpredictability of maternal mood (*r* = .17), as well as predictive validity for risk of depression (*r* = 0.42), anhedonia (*r* = 0.37), and anxiety (*r* = 0.43) (16, 17, 22).

### Measurement of BOLD Response to Novel Relative to Familiar Stimuli

The neural response to novel relative to familiar scenes was characterized using a task of incidental memory encoding (Figure 2). This task was designed to be child friendly and consisted of landscape pictures (selected from the National Geographic library) that were either “familiar” (seen in a training period before the fMRI session) or “novel” (not seen in the training period). Children were instructed to press the left (index finger, dominant hand) button when an animal was in the scene, and the right (middle finger, dominant hand) button when no animal was present. An additional “baseline” task was included that required the participants to discriminate between two noise boxes of differing brightness against a noise background. The intensity of the boxes was randomized, and the participant was instructed to press the button matching the brightest opacity (left or right). Four sets of three 30s block types were presented in random order and counterbalanced across participants. Within blocks, landscape images (12 per block) were presented for 2.5s. The primary contrast of interest for this study was novel minus familiar trials. We also considered novel minus baseline and familiar minus baseline trials in follow-up analyses.

**Figure 2.**
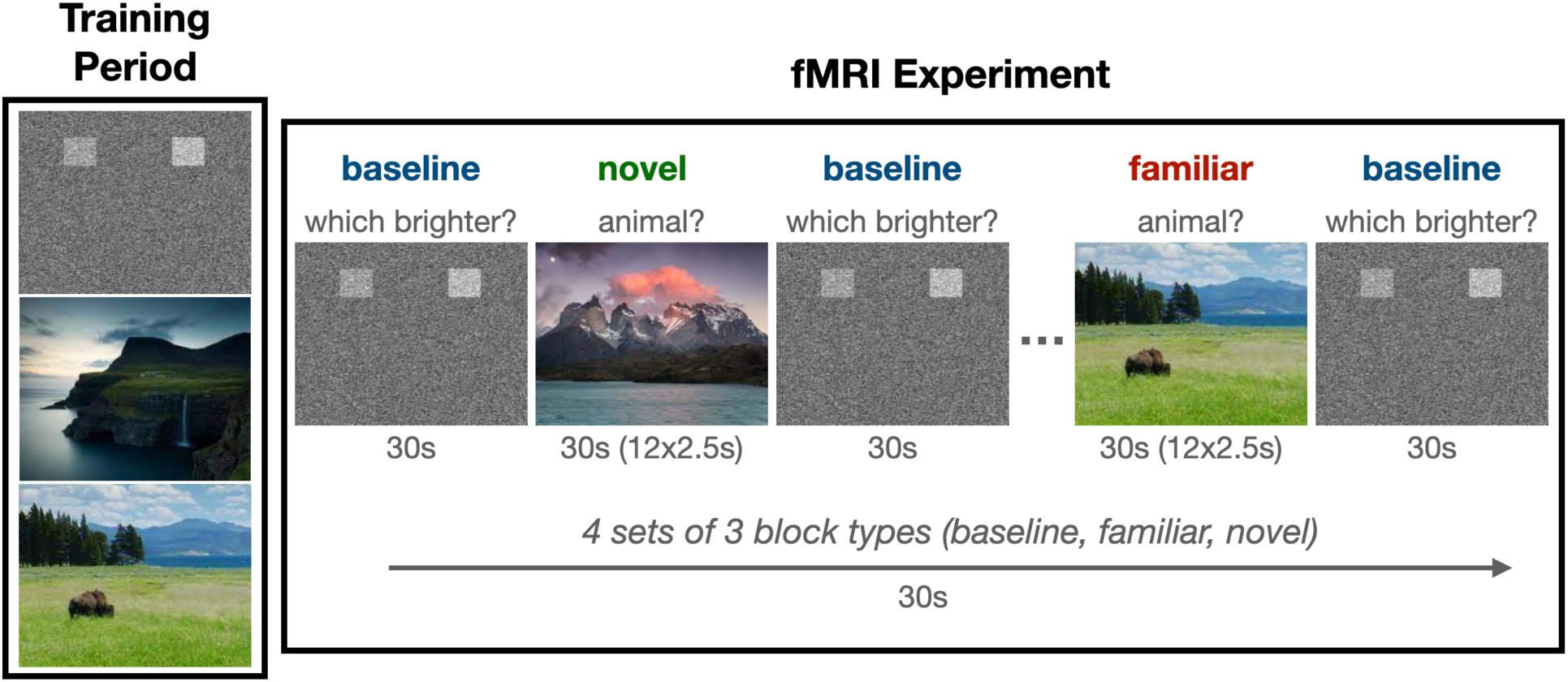
Task Paradigm. Participants underwent an initial training session prior to imaging to familiarize themselves with the task and outdoor scenes. Novel and familiar outdoor scenes were presented in a block design.

### MR Image Acquisition and Processing

Whole-brain MRI was performed using a Philips 3T MRI scanner equipped with a 32-channel head coil. Participants were instructed to remain awake throughout the imaging protocol. T1-weighted 3D MPRAGE (SENSE factor =2.4, 1mm iso, TR/TE = 8/3.7 ms, matrix=256×204, 208 slices, FA=8°, scan duration of 9m 46s) images were acquired for anatomical reference. BOLD-weighted images were acquired using a single shot interleaved EPI sequence (flip angle =64°, TR=2.5s, ∼3mm iso, TE=30ms, matrix=64×61, 51 slices, SENSE reduction factor of 2.5, scan duration of 6m 40s).

EPI timeseries preprocessing was consistent with our previous work (41), utilizing a combination of FMRIPrep (version 20.1.1) (42, 43) and XCP engine (44) imaging pipelines (version 1.2.1). First, T1-weighted images were corrected for b0 inhomogeneities (N4BiasFieldCorrection), skull-stripped (antsBrainExtraction), classified into tissue type (gray matter, white matter, CSF; FSL-FAST), put into template space (ICBM 152 nonlinear asymmetric; antsRegistration), and used as a reference throughout the fMRI preprocessing pipeline. Distortion correction was estimated using ANTS’ symmetrical normalization tool and FSL-MCFLIRT was used for co-registration. Finally, ICA-based automatic removal of motion artifacts was performed on the normalized and spatially smoothed (6mm Gaussian) BOLD time series, and a bandpass (0.01-0.08Hz Butterworth) filter was applied for signal detrending and high-frequency noise removal.

Image post-processing was performed using FSL (FMRIB’s Software Library, www.fmrib.ox.ac.uk/fsl) (45). Specifically, the functional data were analyzed within the framework of the General Linear Model using FEAT (FMRI Expert Analysis Tool, Version 6.00). Three explanatory variables (EVs), corresponding to the conditions ‘novel’, ‘familiar’, and ‘baseline’, were modeled. Each EV was convolved with a double-gamma hemodynamic response function to generate the model time courses and fit to the fMRI data. Parameter estimates (z-scores) for the contrasts of interest (primary outcome: novel vs. familiar; sensitivity analyses: novel vs. baseline, and familiar vs. baseline) were derived on a voxelwise basis.

### MR Image Feature Extraction

An *a priori* hypothesis-based analytical approach was used to focus on regional limbic activation while improving sensitivity and limiting false positive detection. Regions-of-interest (ROIs) were limited to the amygdala and hippocampus. We divided the hippocampal ROI into ventral (body and tail) and dorsal (head) for both empirical and conceptual considerations. Empirically, head and tail activation are not well correlated. Conceptually, the spatial heterogeneity of hippocampal regions is recognized as important in episodic memory versus emotional memory. ROIs were defined based on the Harvard-Oxford atlas(46) and used to extract regional means for further analysis.

A secondary outcome was derived based on sex-specific findings suggesting that early life adversity is related to the amygdala response to novel stimuli. Specifically, Psychophysiological Interaction (PPI) parameter estimates were derived that are reflective of the degree to which the right amygdala integrates (signal correlation) with the other ROIs based on task condition. This model was specified in FSL’s FEAT using main effects of task condition and right amygdala signal, and the multiplicative factor (interaction) of the two (*i.e.,* signal correlation conditioned on task condition). ROIs were defined as above and used to extract regional values for further analysis. Collectively, the above analyses were designed to capture two different but related constructs of function: 1) traditional amygdala/hippocampal response to task conditions, and 2) task condition-dependent integration/correlation with the right amygdala.

### Statistical Modeling Approach

Preliminary analyses were conducted using parameter estimates to characterize baseline activation (novel vs. familiar) and its association with age and sex. Three independent mixed-effects regression models were developed, each for a different ROI. P-values reported throughout are uncorrected for multiple comparisons due to the relatively small number of comparisons in the primary hypothesis-based analyses (three) and/or secondary nature of follow-up analyses (potential confounding covariates and psychophysiological interaction). The models were formulated to predict the bilateral average of each ROI as a function of age and sex, with random intercepts for subjects to account for intra-subject variability. All N=168 unique individuals with a total of 241 visits were considered for preliminary analysis to provide the most precise estimate of age and sex effects possible.

“Primary” analyses considered sex-specific activation (novel vs. familiar) in the context of inter-individual variation in early life unpredictability (QUIC). Parsimonious mixed effects models (main effects of QUIC, age, and sex, with random intercepts for subjects) were developed to identify the main effect of QUIC on activation (novel vs. familiar), followed by repeating this model with an additional QUIC by sex interaction term. Models stratified by sex were used to help interpret the interaction terms. While these analyses provided initial insights, further sensitivity analyses were conducted to test rigorously their robustness and to extend the findings. Specifically, we considered: 1) lateralized effects (six total models), 2) sensitivity to how age was modeled (oldest visit only, aggregating activation across visits, and limiting age to a narrow band [10-13 years]), 3) potentially confounding factors (negative life events and income-to-needs ratio), and 4) task conditions relative to baseline (i.e., novel vs. baseline and familiar vs. baseline). N=150 individuals with a total of 223 visits were available for primary analysis that included QUIC measurement.

“Secondary” analyses using PPI measures were conducted to further extend the findings with respect to task condition-dependent integration with the right amygdala. Specifically, the above models (main effect of QUIC, and QUIC by sex interaction) were repeated using the PPI with the right amygdala (five models, one for each lateral ROI) as the outcome variable.

### Covariates

Sex was determined based on biological assignment at birth and gender was self-reported. It should be noted that one participant reported a gender different from their assignment at birth. This individual was considered in a sensitivity analysis to determine the effect of using self-reported gender rather than biological assignment at birth. The analysis showed minimal effect, so detailed results concerning this case are not discussed further in this report. Income-to-needs ratios (INR) and childhood experiences of negative life events were assessed via self-report completed at the time of QUIC assessment and used here to test the association with unpredictability beyond established ACEs such as for socioeconomic status and negative life events (47), respectively. To account for a total of n=5 missing INR values, we conducted a multiple imputation analysis using the “mice” package in R (20 imputations, predictive mean matching).

## Results

### Novel vs. Familiar BOLD Contrast Is Independent of Sex and of Age at MRI

Significant group activation (novel vs. familiar contrast) was observed in the hippocampal head (t=-1.5, p<0.001), but not the tail of the hippocampus or amygdala (p>0.1). The direction of effect suggests greater activation in the head of the hippocampus when presented with familiar relative to novel stimuli. The novel relative to familiar BOLD contrast was not associated with age, sex, or age in a sex-specific manner (*i.e.,* age by sex interaction, all p’s>0.1) in any of the three bilateral ROIs.

### Hippocampus and Amygdala Novelty Signals Associated with Early Life Unpredictability in a Sex-Specific Manner

No main effect of QUIC was observed on the novel relative to familiar BOLD contrast (p>0.1). Notably, there was a significant QUIC by sex interaction (Figure 3) in the head of the hippocampus (t=2.0, p=0.047) and amygdala (t=2.7, p=0.007). Specifically, the novel vs. familiar BOLD contrast negatively associated with QUIC scores in males (t_headHipp_=-2.0, p_headHipp_=0.045; t_amyg_=-2.8, p_amyg_=0.007). In contrast, the effect was not present in females (p>0.1). Additional analyses suggested that these associations were specific to unpredictability: they largely persist over and above potentially confounding factors reflecting traditional measures (negative life events and INR) of ELA (confound-adjusted QUIC by sex statistics without imputation: n=147; t_amyg_=2.6, p_amyg_=0.009; t_headHipp_=1.9, p_headHipp_=0.062; pooled confidence intervals with imputation: n=150; β_amyg_=0.022±0.010 (S.E.) 95% CI [0.006 0.047]; β_headHipp_=0.018±0.009 (S.E.) 95% CI [-0.001 0.037]). Sensitivity analyses provided two additional observations: 1) A left-right asymmetry (see Supplementary Table S1) suggests a larger effect in the right amygdala, which prompted focusing on the right amygdala for secondary PPI analyses, and 2) sex-specific amygdala associations are robust to modeling and sample selection with respect to age at visit (see Supplementary Table S2). In addition, the observed effects endured when considering novel relative to baseline conditions in the amygdala (task-specific QUIC by sex statistics: t_amyg,novel_=2.0, p_amyg,novel_=0.047), but not familiar relative to baseline conditions (p>0.1). Collectively, these findings provide robust evidence that early life unpredictability is negatively associated with the neural response of key limbic structures to novel scenes in a sex-specific manner.

**Figure 3.**
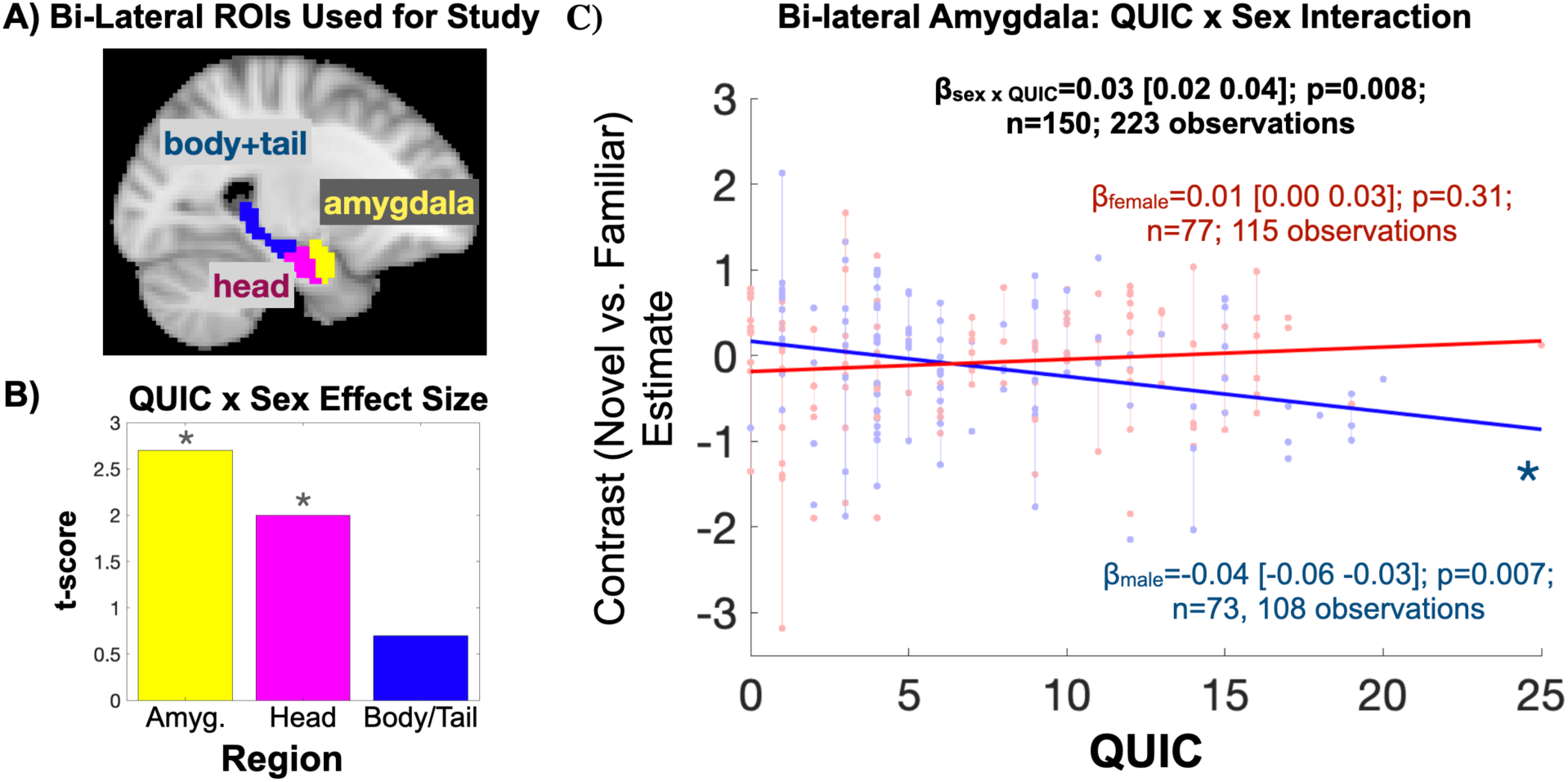
Early Life Unpredictability (QUIC) Is Associated with Amygdala and Hippocampal Head Activation to Novel vs. Familiar Scenes in a Sex-Specific Manner. A) The amygdala and hippocampus (split by head and body/tail) were considered for analysis. B) The amygdala and head of the hippocampus functional BOLD activation associate with QUIC scores in a sex-specific manner (* denotes p<0.05). C) Scatter plot depicting sex-specific associations. In stratified analyses, QUIC scores were associated with amygdala and head of hippocampus (not shown) activation in males, but not females.

### Amygdala-Hippocampal Connectivity is Associated with Early Life Unpredictability in a Sex– and Condition-Specific Manner

Based on the above findings, psychophysiological interaction (PPI) analysis was performed to probe the sex-specific associations between QUIC and right amygdala connectivity to its lateral analogue and the left/right head/tail of the hippocampus in a condition-specific manner. No main effects of QUIC on PPI measures were observed. A significant QUIC by sex interaction was observed in the context of condition-specific connectivity between the right amygdala and the left amygdala, as well as between the right amygdala and the left/right head and tail of the hippocampus (Figure 4). Stratified analyses provided evidence that the association between QUIC and condition-specific connectivity with the right amygdala is stronger in females. Broadly, these findings suggest that functional connectivity to the amygdala in males exposed to early childhood unpredictability is relatively greater in the novel condition, whereas in females, functional connectivity to the amygdala is relatively greater in the familiar condition. Thus, this finding supports the idea that early life unpredictability is associated with sex-specific patterns of limbic connectivity, with males showing a bias towards novel stimuli and females towards familiar stimuli.

**Figure 4.**
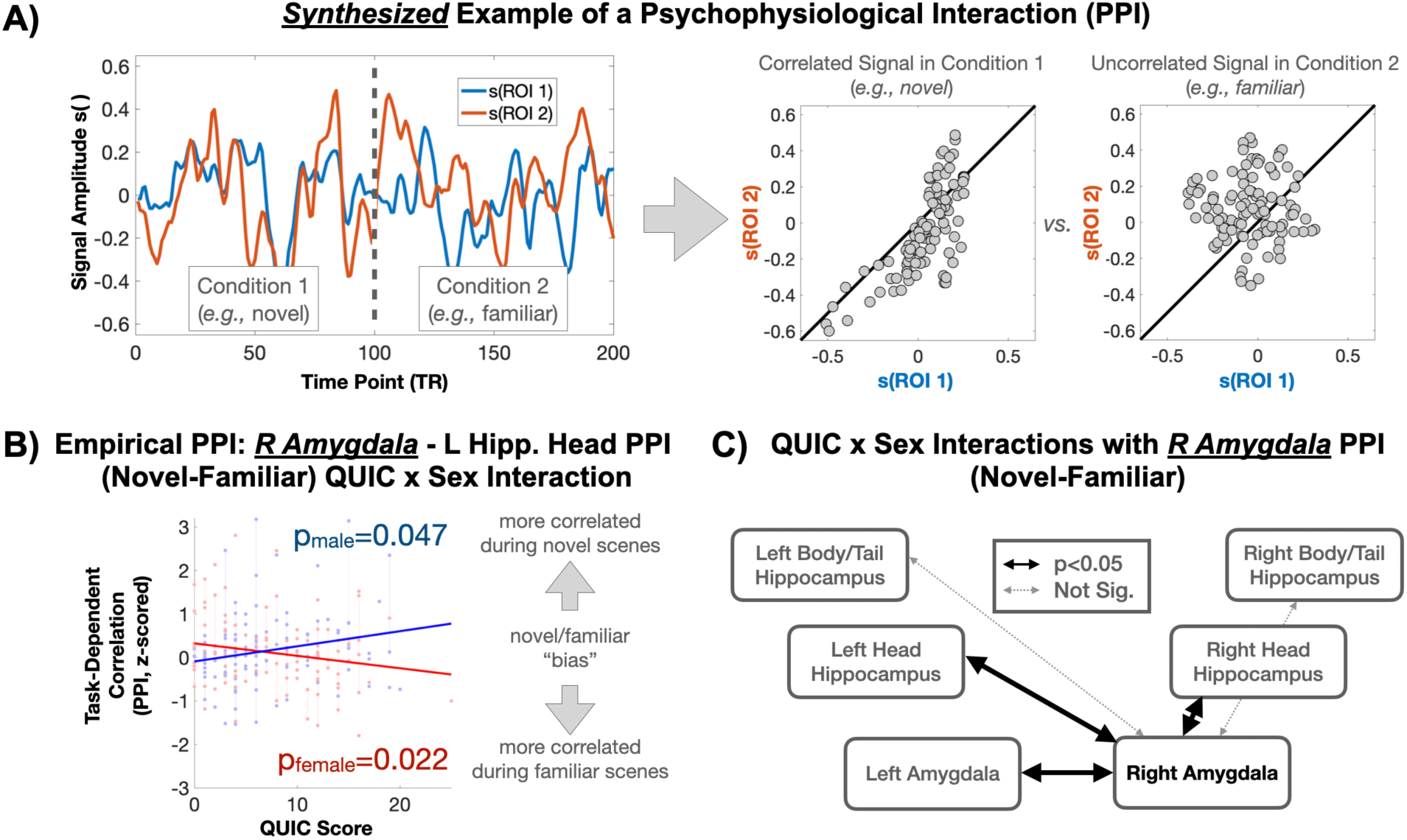
Early Life Unpredictability Is Associated with Task-Based Functional Connectivity of the Right Amygdala in a Sex-Specific Manner. A) Synthesized examples of raw signals occurring in a Psychophysiological Interaction (PPI, upper left panel) and their task-dependent correlation structure (upper right panel). B) Empirical scatter plot showing sex-specific associations between early-life unpredictability (QUIC) and PPI-based task-dependent correlations. In a sex-stratified analysis, QUIC scores were significantly (p<0.05) associated with PPI measures in both sexes but in opposing directions. C) QUIC by sex interactions are depicted across the observed amygdala and hippocampus ROIs.

## Discussion

This study provides new evidence that exposure to early life unpredictability has sex-specific implications for the functional organization of limbic circuits and their responses to novelty. Specifically, we observed sex-specific associations between early life unpredictability and limbic regional (hippocampus and amygdala) BOLD activation/connectivity in response to novel relative to familiar scenes. In males, we observed that higher early life unpredictability was associated with decreased limbic BOLD *activation* in response to novel stimuli. In females, higher unpredictability was associated with decreased condition-specific (novel relative to familiar) functional *connectivity* to the amygdala, while the opposite pattern was observed in males. Taken together, these findings suggest that early life unpredictability has ramifications for both male and female limbic system development yet manifests in different ways. Importantly, the observed associations were not explained by more commonly studied forms of ELA such as socioeconomic disadvantage and negative life events. This work adds to a growing body of evidence suggesting that the effects of early life unpredictability on neurodevelopment may be distinct from those reflective of overall exposure to adversity.

This study employed a hypothesis-driven approach designed to understand the effects of early life unpredictability on functional activation and connectivity within and between key brain regions involved in learning novel experiences, specifically the amygdala and hippocampus. We posited that exposure to unpredictable signals from the parents and environment during a sensitive developmental period may alter processing of neural responses to novelty later in life. Consistent with this concept, we found that the amygdala and hippocampus are functionally altered in response to early life unpredictability.

The amygdala has a well-established role in emotional processing (48), motivating its widespread use as a target for understanding the putative pathway between ELA and mental health outcomes (49–52). Notably, the amygdala asymmetry observed in our study aligns with previous findings (53). Our results join a diverse body of work demonstrating a spectrum of changes in amygdala activation and connectivity in response to various forms of ELA. Interestingly, the sex-specific plasticity we observed in response to early-life unpredictability extends previously reported non-sex-specific findings of altered fronto-amygdala connectivity in cases of neglect or maltreatment (54). This suggests that different types of early-life adversity may distinctly shape amygdala function, likely through unique neurodevelopmental mechanisms.

In the hippocampus, we observed a significant functional distinction between the head (anterior) and the body/tail (posterior) regions, particularly in response to novel stimuli. The head also showed more pronounced condition-specific integration with the amygdala, a key center for emotional processing. These findings are consistent with the anterior-posterior emotion-cognition segregation hypothesis (55–57), which posits that the head of the hippocampus is differentially involved in emotional processing relative to the body/tail region. While this anterior-posterior functional gradation is more pronounced in rodent models and remains debated in humans, our findings of increased gradation in individuals exposed to unpredictable early life environments suggest that this gradient may become more distinct following developmental insults. One complimentary interpretation of these findings is that the gradation observed here simply reflects a greater integration between the anterior hippocampus and the amygdala, rather than a functional division of labor suggested by the anterior-posterior emotion-cognition segregation hypothesis.

We report significant and opposing effects of unpredictability in males and females, highlighting sex differences in the processing of novel stimuli. Our observations suggest that early life unpredictability influences limbic circuitry activation and connectivity, leading to a bias towards processing either novel or familiar stimuli, with this bias differing between males and females in response to ELA. These findings may have implications for how individuals process information in novel (*e.g.,* first day of school) versus familiar (*e.g.,* end of the school year) situations, with sex-specific impacts of ELA potentially contributing to differences in mental health disorders (*e.g.,* anxiety and/or withdrawn behaviors) characterized by novelty avoidance. Notably, these concepts are consistent with an emerging body of preclinical evidence illustrating distinct phenotypic consequences of early life unpredictability in male and female rodents. For example, early life unpredictability has been experimentally shown to increase *anhedonic* behavior in males, while unpredictable signals have been linked to increases in *hedonic* (e.g., drug-seeking) behaviors in females (11, 58–60). Thus, these findings highlight the importance of considering sex differences in the study of ELA and its long-term effects on mental health (1, 7, 33).

The current study’s limitations should be carefully considered. We did not include behavioral outcomes such as novelty-seeking, internalizing, or risk-taking behaviors. Although we contextualize our findings within the existing literature, further research is necessary to directly link the observed neural patterns with their behavioral relevance and potential mediating roles. Additionally, condition-specific approaches to characterizing functional connectivity have a demonstrated potential to induce spurious findings as they rely on precise specification of task-based activation for nuisance regression (61). While we cannot entirely rule this out, the choice of block-designed stimuli minimizes misspecification errors associated with onset and cessation of stimuli and, furthermore, the associations with unpredictable early life experiences are unlikely to be confounded in this case. Finally, while the current sample and study design may be sufficiently powered, the typically small magnitude of brain-behavior effects suggests the need for additional future studies. Notably, the upcoming HBCD study, which will include a short-form QUIC concurrent with early life MRI assessments from birth to six-years age (62), provides an opportunity to generalize the current work and to explore the timing and onset of effects, addressing a current “limitation” where brain measures were collected long after exposure to the unpredictability measure of the child’s environment.

In summary, our findings demonstrate that unpredictability in early life differentially impacts the functional activation and connectivity of key limbic regions—amygdala and hippocampus—in the context of novel vs. familiar stimuli and that these effects are sex-dependent. Importantly, we examine a novel form of adversity (unpredictable signals), which exerts enduring consequences beyond those established for other dimensions of ELA. By identifying unpredictable signals as a distinct form of adversity with lasting neurodevelopmental consequences, our study further highlights the critical importance of stable early environments for healthy brain development (11, 63).

## Acknowledgements

The authors thank the families who participated in these projects. We also thank the dedicated staff at the Early Human and Lifespan Development Program and the Women and Children’s Health and Well-Being project. This research was supported by grants from the National Institutes of Health (P50MH96889 R01NS41298, R01HD28413, R01HD51852, R01NS41298, R00HD100593, R01 HL155744, R01 MH109662) and VA (BX006186).

