## supplement for "Sex-Specific Effects of Early Life Unpredictability on Hippocampal and Amygdala Responses to Novelty in Adolescents"

#### *Supplementary Materials*

### Supplement S1: Evidence for Rightward Asymmetry in Sex-Specific Associations with QUIC

Bi-lateral ROIs were considered in the main manuscript to reduce comparisons and increase sensitivity. However, based on a strong body of literature demonstrating laterality of developmental programming effects in limbic region outcomes(1), we conducted follow-up sensitivity analyses splitting ROIs into their lateral components. Six independent linear mixed effects models (N=150 unique individuals, 223 observations) were tested consistent with those used in the main manuscript. We observed the strongest sex-specific effects in the right hemisphere and bi-laterally in the amygdala (Supplementary Table 1). Based on: 1) pre-existing evidence for asymmetry, 2) the strongest effects having been observed in the amygdala, and 3) the premise that bi-lateral averaging of spatially separated time courses may obfuscate important frequency characteristics used for connectivity measurement, the right amygdala was chosen for secondary Psychophysiological Interaction (PPI) analyses targeted towards understanding condition-specific signal correlations within a lateral limbic network (amygdala, hippocampal head, hippocampal tail/body).

| | **Left** | | **Right** | | **Bi-lateral** | |
|---|---|---|---|---|---|---|
| | **t** | **p** | **t** | **p** | **t** | **p** |
| **Amygdala** | *2.24* | *0.026* | *2.42* | *0.015* | *2.69* | *0.007* |
| **Hippocampal Head** | 1.44 | 0.152 | *2.19* | *0.029* | *2.00* | *0.047* |
| **Hippocampal Body/Tail** | 0.44 | 0.657 | 1.02 | 0.310 | 0.74 | 0.460 |

**Supplementary Table 1. Lateralized Model Statistics.** Six additional independent models were conducted to test for a laterality of effects. Evidence supports a rightward asymmetry and localizes the amygdala as having the strongest effect. Bi-lateral statistics (right columns) are redundant with those reported in the main manuscript but provided for reference.

### Supplement S2: Sex-Specific QUIC Associations in the Amygdala Are Robust to Age Distribution and Modeling Considerations in the Context of Participant Age

The main manuscript used a mixed-effects model to account for sampling heterogeneity and spread in ages at visit. Given that late childhood/early adolescence is a period of rapid development, we sought to understand the influence of sampling and modeling on model statistics. Specifically, we considered three additional approaches: 1) considering the oldest visit only, 2) aggregating (averaging age and fMRI BOLD contrast) across visits, and 3) repeating analyses in a narrower age band (10-13 years old). Broadly, we found that the primary sex-specific associations with QUIC are robust to age considerations in the amygdala. Specifically, this is demonstrated through continued statistical significance (despite the reduced sample size) and/or a persistent effect size and direction.

| | **n (observations)** | **t QUIC x Sex** | **p QUIC x Sex** |
|---|---|---|---|
| *Bi-lateral Amygdala* | | | |
| **Reference** | n=150 (223) | *2.69* | *0.007* |
| **Oldest Visit** | n=150 (150) | 1.76 | 0.080 |
| **Averaged** | n=150 (150) | *2.42* | *0.017* |
| **Narrow Range** | n=105 (107) | *2.06* | *0.041* |
| *Bi-lateral Hippocampal Head* | | | |
| **Reference** | n=150 (223) | *2.00* | *0.047* |
| **Oldest Visit** | n=150 (150) | 1.56 | 0.12 |
| **Averaged** | n=150 (150) | 1.81 | 0.07 |
| **Narrow Range** | n=105 (107) | 1.12 | 0.27 |
| *Bi-lateral Hippocampal Body/Tail* | | | |
| **Reference** | n=150 (223) | 0.74 | 0.460 |
| **Oldest Visit** | n=150 (150) | 1.01 | 0.315 |
| **Averaged** | n=150 (150) | 0.61 | 0.544 |
| **Narrow Range** | n=105 (107) | -0.08 | 0.929 |

**Supplementary Table 2. Models Repeated Under Different Sampling and Modeling Strategies.** Primary sex-specific associations with QUIC were largely robust to sampling and modeling strategies in the context of participant age.

1. Teicher MH, Khan A. Childhood Maltreatment, Cortical and Amygdala Morphometry, Functional Connectivity, Laterality, and Psychopathology. Child Maltreat. 2019;24(4):458-65.
